# Reversal of β-amyloid induced microglial toxicity *in vitro* by activation of Fpr2/3

**DOI:** 10.1101/2020.02.13.947051

**Authors:** Edward S. Wickstead, Husnain A. Karim, Roberta E. Manuel, Christopher Biggs, Stephen J. Getting, Simon McArthur

## Abstract

**Background and Purpose:** Microglial inflammatory activity is thought to be a major contributor to the pathology of neurodegenerative conditions such as Alzheimer’s disease (AD), and strategies to restrain their behaviour are under active investigation. Classically, anti-inflammatory approaches aim to suppress pro-inflammatory mediator production, but exploitation of inflammatory resolution, the endogenous process whereby an inflammatory reaction is terminated, has not been fully investigated as a therapeutic approach in AD. In this study, we sought to provide proof-of-principal that the major pro-resolving actor, formyl peptide receptor 2, Fpr2, could be targeted to reverse microglial activation induced by the AD-associated pro-inflammatory stimulus, oligomeric β-amyloid (oAβ).

**Experimental Approach:** The immortalised murine microglial cell line BV2 was employed as a model system to investigate the pro-resolving effects of the Fpr2 ligand QC1 upon oAβ-induced inflammatory, oxidative and metabolic behaviour. Cytotoxic behaviour of BV2 cells was assessed through use of co-cultures with retinoic acid-differentiated human SH-SY5Y cells.

**Key Results:** Stimulation of BV2 cells with oAβ at 100nM did not induce classical inflammatory marker production but did stimulate production of reactive oxygen species (ROS), an effect that could be reversed by subsequent treatment with the Fpr2 ligand QC1. Further investigation revealed that oAβ-induced ROS production was associated with NADPH oxidase activation and a shift in BV2 cell metabolic phenotype, activating the pentose phosphate pathway and NADPH production, changes that were again reversed by QC1 treatment. Microglial oAβ-stimulated ROS production was sufficient to induce apoptosis of bystander SH-SY5Y cells, an effect that could be prevented by QC1 treatment.

**Conclusion and Implications:** In this study, we provide proof-of-concept data that indicate exploitation of the pro-resolving receptor Fpr2 can reverse damaging oAβ-induced microglial activation. Future strategies aiming to restrain neuroinflammation in conditions such as AD should examine pro-resolving actors as a mechanism to harness the brain’s endogenous healing pathways and limit neuroinflammatory damage.

## Background

AD is the single greatest cause of dementia, affecting approximately 4% of individuals aged over 65 years and with a global disease burden of around 37 million individuals [1]. This figure is set to increase as the population ages, and is expected to reach around 78 million people by 2050 [2]. There are currently no effective treatments for the condition.

Whilst the two core pathological lesions of AD, extracellular β-amyloid (Aβ) plaques and intraneuronal tau tangles, have long been studied, the contribution to pathology provided by neuroinflammation, and the role of the microglia in AD pathogenesis, has only recently been appreciated [3,4]. Several lines of evidence indicate a pathological role for microglial activity: studies of genetic risk factors for idiopathic AD have identified numerous immune-related risk loci, clinical imaging studies indicate a positive correlation between microglial activity and both Aβ load and neurodegeneration [5], and chronic neuroinflammation is a feature of multiple independent animal models of the disease [6]. More directly, Aβ can act as a damage-associated molecular pattern [7], stimulating microglial activation through a range of different receptors, including the receptor for advanced glycation end products, toll-like receptors, and CD36 [8].

Under normal conditions, inflammation is self-resolving, with numerous factors acting to ‘switch off’ inflammatory processes [9]. A central actor in this process is the G protein-coupled receptor formyl peptide receptor 2 (FPR2) or its murine functional homologues Fpr2/3 [10]. Strong evidence exists for the pro-resolving potential of this receptor in peripheral inflammation, where it promotes neutrophil apoptosis [11], and regulates monocyte/macrophage recruitment [12,13], phenotype [14] and behaviour [15]. Importantly, protective effects have been identified for this receptor in diverse inflammatory settings, including sepsis [16], heart failure [17] and atherosclerosis [18].

Expression of FPR2 within the brain has been reported in the endothelium and in selected hippocampal and cerebellar neurones [19], but it is also expressed by microglia [20], and is rapidly upregulated following inflammatory insult [21]. Significantly, FPR2 expression has been reported in inflammatory cells infiltrating Aβ plaques in AD [22], is involved in chemotaxis to high concentrations of Aβ [23] and has been indirectly implicated in microglial Aβ phagocytosis [24]. Given the importance of this receptor in the resolution of peripheral inflammation, we hypothesised that FPR2 agonists would be able to reverse the pro-inflammatory effects of Aβ upon microglia, restoring normal homeostasis.

## Methods

### Drugs & Reagents

The FPR2 agonist Quin-C1 (QC1; 4-Butoxy-N-[1,4-dihydro-2-(4-methoxyphenyl)-4-oxo-3(2H)-quinazolinyl]benzamide) and antagonist WRW_4_ (Trp-Arg-Trp-Trp-Trp-Trp-NH_2_) were purchased from Tocris Ltd, UK. Isolated and purified lipopolysaccharides developed in *Escherichia* coli, serotype O111:B4 were purchased from Perck Millipore, Ltd, UK. HFIP-treated human Aβ_1-42_ peptide was purchased from JPT Peptide Technologies, Berlin, Germany.

### Aβ oligomerisation

HFIP-treated Aβ_1-42_ stored at −80°C in DMSO was oligomerised by dilution and vortexing in PBS followed by incubation overnight at 4°C [25]. Oligomer formation was confirmed by native Tricine-SDS-polyacrylamide gel electrophoresis. Briefly, 2 μg oligomeric Aβ (oAβ) was resuspended in non-denaturing sample buffer (62.5 mM Tris-base, 25% glycerol, 1% (w/v) Coomassie Blue R-250) and loaded onto a 10% acrylamide:bis-acrylamide gel and separated by electrophoresis alongside molecular weight markers. Gels were incubated with Coomassie stain (60mg/l Coomassie Blue R-250, 10% v/v acetic acid, both Sigma, UK) Following 24h de-staining in 10% v/v acetic acid, 50% v/v methanol (Sigma, UK), gels were imaged using a ChemiDoc MP Imaging System (Bio-Rad Ltd., UK). Oligomeric Aβ migrated at approximately 35kDa, indicating the presence of hexamers/heptamers (Supplemental Figure 1).

### Cell culture

The murine microglial line BV2 were a generous gift from Prof. E. Blasi (Università degli Studi di Modena e Reggio Emilia, Italy); the human neuroblastoma SH-SY5Y line was purchased from the European Collection of Authenticated Cell Cultures (ECACC, Salisbury, UK). Both lines were cultured in DMEM medium supplemented with 5% fetal calf serum and 100 μM non-essential amino acids, 2 mM L-alanyl-L-glutamine and 50 mg/ml penicillin-streptomycin (all Thermofisher Scientific, UK) at 37°C in a 5% CO_2_ atmosphere. SH-SY5Y cells were differentiated to a neurone-like phenotype prior to experimentation by incubation with 10 μM trans-retinoic acid (Sigma, UK) for 5 days [26].

### Reactive oxygen species (ROS) assays

Total intracellular ROS production was quantified using 6-chloromethyl-2’,7’-dichlorodihydrofluorescein diacetate, acetyl ester (CM-H_2_DCFDA; Thermofisher Scientific, UK) according to the manufacturer’s recommendations. Briefly, cells were plated at 200,000 cells/cm^2^ in phenol red-free DMEM, serum starved overnight and pre-loaded with 5 μM CM-H_2_DCFDA for 20 minutes at 37°C. Following removal of unbound dye, fresh phenol red free-DMEM was added and experimental treatments were begun. Following administration of treatments, cellular fluorescence was determined every 5 minutes for 1 hr at 37°C using a CLARIOstar fluorescence microplate reader (BMG Labtech, Germany) with excitation and emission filters set at 492nm and 517nm respectively.

Mitochondrial superoxide production was quantified using the tracer MitoSOX Red (Thermofisher Scientific, UK) according to the manufacturer’s recommendations and a loading concentration of 2.5 μM. Following administration of treatments, cellular fluorescence was determined every 5 minutes for 1 hr at 37°C using a CLARIOstar fluorescence microplate reader (BMG Labtech, Germany) with excitation and emission filters set at 510nm and 580nm respectively.

Hydrogen peroxide production was quantified using the ROS-Glo H_2_O_2_ assay (Promega, Southampton, UK) according to the manufacturer’s recommendations. Following experimental treatment, luminescence of cell lysates at 37°C was determined using a CLARIOstar luminescence microplate reader (BMG Labtech, Germany), in comparison to a H_2_O_2_ standard curve (0.013 μM – 10 mM).

### GSH:GSSG ratio analysis

The ratio of reduced (GSH) to oxidised (GSSG) glutathione was determined using a commercial assay (GSH:GSSG-Glo assay, Promega Co, Southampton, UK) according to the manufacturer’s instructions, with cells plated at 200,000 cells/cm^2^ on black walled 96-well plates. A CLARIOstar spectrophotometer (BMG Labtech, Germany) was used to measure relative luminescence with comparison to a total glutathione standard curve (0.25 μM – 16 μM).

### Cytokine ELISA

Tumour necrosis factor alpha (TNFα), was assayed by murine-specific sandwich ELISA using commercially available kits, according to the manufacturer’s protocols (ThermoFisher Scientific, UK). A CLARIOstar spectrophotometer (BMG Labtech, Germany) was used to measure absorbance at 450 nm.

### E. coli bioparticle phagocytosis

Microglial phagocytic capacity was determined using BODIPY-FL conjugated *Escherichia coli* (K-12 strain) bioparticles (ThermoFisher Scientific, UK). Following experimental treatments, cells were incubated with bioparticle conjugates at a ratio of 50 particles per cell in PBS for 30 minutes at 37°C in the dark. Cells were washed, fluorescence of non-engulfed particles was quenched by addition of 0.2% Trypan blue (ThermoFisher Scientific, UK) for 1 min, and cellular fluorescence was determined using a FACS Canto II flow cytometer (BD Biosciences, UK) equipped with a 488nm laser and FlowJo 8.8.1 software (Treestar Inc. FL, USA). A total of 10,000 singlet events per sample were quantified.

### Flow cytometry

BV2 or SH-SY5Y cells alone or in co-culture were labelled with APC-conjugated rat monoclonal anti-mouse CD11b, PE-Cy7-conjugated rat monoclonal anti-mouse CD40 (Biolegend, UK) or PerCP-Cy5.5-conjugated mouse monoclonal anti-human CD200 (all Biolegend, UK) for analysis by flow cytometry. Immunofluorescence was analysed for 10,000 singlet events per sample using a BD FACSCanto II (BD Biosciences, UK) flow cytometer; data were analysed using FlowJo 8.8.1 software (Treestar Inc., CA, USA).

### Annexin A5 apoptosis assay

SH-SY5Y cells were differentiated as described above and treated according to experimental design, either alone or in co-culture with BV2 cells. cultures were in PBS, detached using a cell scraper and incubated with FITC-conjugated annexin A5 (0.45 μg/ml in 0.01 M PBS, 0.1% bovine serum albumin, 1 mM CaCl_2_), and in the case of co-cultures, APC-conjugated rat monoclonal anti-mouse CD11b and PerCP-Cy5.5-conjugated mouse monoclonal anti-human CD200 (all Biolegend, UK) on ice in the dark for 30 min. Samples were washed and analysed by flow cytometry. Immunofluorescence was analysed for 10,000 singlet events per sample using a BD FACSCanto II (BD Biosciences, UK) flow cytometer; data were analysed using FlowJo 8.8.1 software (Treestar Inc., CA, USA).

### Western blot analysis

Samples boiled in 6× Laemmli buffer were subjected to standard SDS-PAGE (10%) and electrophoretically blotted onto Immobilon-P polyvinylidene difluoride membranes (Merck, UK). Total protein was quantified using Ponceau S staining (Merck, UK) and membranes were blotted using antibodies raised against murine haem oxygenase-1 (HO-1; rabbit polyclonal, 1:1000, Cell Signaling Technology, Leiden, The Netherlands) or superoxide dismutase 2 (SOD2; rabbit monoclonal, 1:1000, Cell Signaling Technology, Leiden, The Netherlands) in Tris-buffer saline solution containing 0.1% Tween-20 and 5% (w/v) non-fat dry milk overnight at 4°C. Membranes were washed with Tris-buffer saline solution containing 0.1% Tween-20, and incubated with secondary antibody (horseradish peroxidase–conjugated goat anti-rabbit 1:5,000; ThermoFisher Scientific, UK), for 90 min at room temperature. Proteins were then detected using enhanced chemiluminescence detection (2.5 mM luminol, 0.4 mM p-coumaric acid, 7.56 mM H_2_O_2_ in 1 M Tris, pH 8.5) and visualised on X-ray film (Scientific Laboratory Supplies Limited, Nottingham, UK). Films were digitized and analysed using ImageJ 1.51w software (National Institutes of Health).

### Immunofluorescence & confocal microscopy

Following experimental treatment, BV2 cells cultured in chambered microslides were fixed by incubation in 2% formaldehyde in PBS for 10 min at 4°C, washed and non-specific antibody binding was minimised by incubation for 30 min at room temperature in PBS containing 10% FCS and 0.05% Triton X-100 (all Thermofisher Scientific, UK). Cells were then incubated with rabbit anti-mouse p67Phox monoclonal antibody (1:500, clone EPR5064, Abcam Ltd, Cambridge, UK) and mouse anti-mouse gp91phox monoclonal antibody (1:50, clone 53, BD Biosciences, UK) overnight at 4°C in PBS with 1% FCS and 0.05% Triton X-100. Cells were washed and incubated with AF488-conjugated goat anti-mouse and AF647-conjugated goat anti-rabbit secondary antibodies (both 1:500, Thermofisher Scientific, UK) in PBS with 1% FCS and 0.05% Triton X-100 at room temperature for 1 hr. Cells were washed with PBS, nuclei were defined by incubation with 180 nM DAPI in ddH_2_O for 5 min, and cells were mounted under Mowiol mounting solution. Cells were imaged using an LSM710 confocal microscope (Leica, UK) fitted with 405nm, 488nm and 647nm lasers and a 63x oil immersion objective lens (NA 1.4mm, working distance 0.17mm). Images were captured with ZEN Black software (Zeiss, Cambridge, UK) and analysed with ImageJ 1.51w (National Institutes of Health, USA).

### Glucose 6-phosphate dehydrogenase activity assay

Glucose 6-phosphate dehydrogenase (G6PD) activity was assessed using a commercial assay (Cell Signalling Technology, UK) according to the manufacturer’s instructions. Following treatment according to experimental design, cells were lysed by ultrasonication (2 × 20s at 20kHz) in assay lysis buffer (22 mM Tris-HCl, 150 mM NaCl, 1 mM Na_2_EDTA, 1 mM EGTA, 1% Triton X-100, 20 mM sodium pyrophosphate, 25 mM sodium fluoride, 1 mM β-glycerophosphate, 1 mM sodium orthovanadate, 1 μg/ml leupeptin, 1 mM phenylmethane sulfonyl fluoride, pH 7.5, 4°C) using a Soniprep 150 (BMG Labtech, UK), centrifuged at 14000g and 4°C for 10 min, and lysates were collected. Samples were diluted to 0.2 mg/ml in assay buffer, incubated at 37°C for 15 min with assay substrate and fluorescence analysed using a CLARIOstar spectrophotometer (BMG Labtech, Germany) with excitation and emission filters set at 540nm and 590nm respectively.

### Mitochondrial function assay

Mitochondrial function was assessed using a Seahorse XF24 cell MitoStress Test (Agilent Technologies, California, USA) according to the manufacturer’s instructions. BV2 cells plated at 2 × 10^6^ cells/cm^2^ were serum starved overnight and treated according to experimental design. Medium was replaced with Seahorse XF DMEM supplemented with 1 g/L glucose and 1 mM sodium pyruvate, pH 7.4 (Sigma, UK) and cells were incubated at 37°C without CO_2_ for 45 min prior to analysis of oxygen consumption rate (OCR) and extracellular acidification rate (ECAR). Basal respiration was initially determined prior to subsequent serial cellular treatments with 4 μM oligomycin, 0.6 μM FCCP and 1 μM rotenone/antimycin A to measure ATP production, maximal respiratory capacity and non-mitochondrial respiration, respectively. For each treatment, readings were taken in triplicate every 5 min. Cells were lysed in RIPA buffer and protein content was assessed by Bradford’s method for sample normalisation. Rates of glycolytic and oxidative ATP production were then calculated as described in [27].

### Statistical analysis

Sample sizes were calculated to detect differences of 15% or more with a power of 0.85 and α set at 5%, calculations being informed by previously published data [28,29]. All experimental data are presented as mean ± SEM, repeated using a minimum of n = 3 independent culture flasks; assays were performed in triplicate. In all cases, normality of distribution was established using the Shapiro–Wilk test, followed by analysis with two-tailed Student’s t tests to compare two groups or, for multiple comparison analysis, one- or two-way ANOVA followed by Tukey’s HSD *post hoc* test, a p<0.05 was considered statistically significant. All statistical analysis was performed using Graph Pad Prism 8 software (GraphPad Software, CA, USA).

## Results

### AD-relevant concentrations of Aβ do not induce an inflammatory response in BV2 microglia

Whilst many studies have investigated the toxic properties of Aβ, these have in general used micromolar concentrations of the peptide, levels which are unlikely to be achieved until the end stages of AD [30]. We sought to determine the potential of Fpr2/3 as a target to control Aβ-driven inflammation earlier in the disease process when oligomeric Aβ is found in the nanomolar range [30], hence we characterised the inflammatory response of BV2 cells to AD-relevant concentrations of Aβ. Initial studies identified a clear dose-dependent increase in BV2 cell reactive oxygen species (ROS) production upon Aβ stimulation (Figure 1A), with 100 nM Aβ stimulating an approximately 2.5 fold increase; this concentration of Aβ was thus used for further investigation.

**Figure 1:**
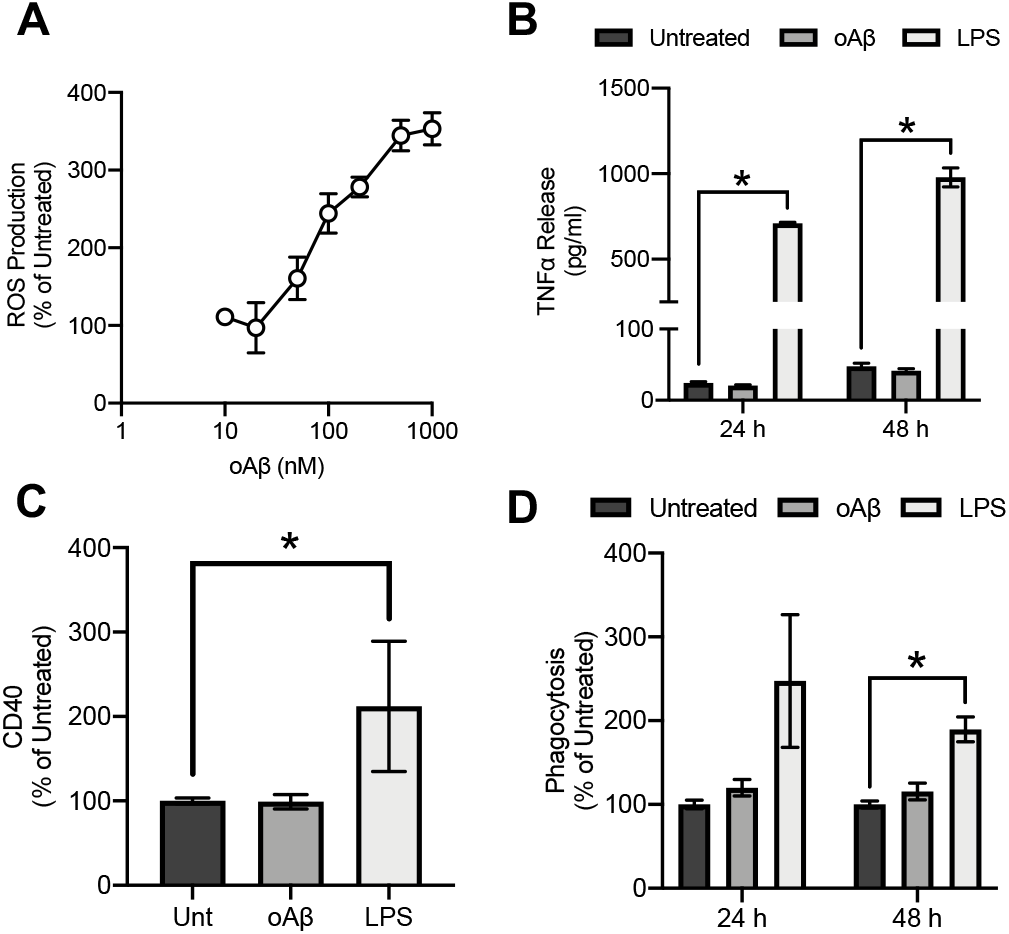
oAβ stimulates microglial ROS production without inducing an inflammatory response. **A)** Treatment with oAβ dose-dependently induces microglial ROS production rate over a 2 hr period. **B)** Treatment of BV2 cells with 50 ng/ml LPS but not 100 nM oAβ increased microglial TNFα production after 24 and 48 hrs exposure. **C)** Treatment for 24 hrs with 50 ng/ml LPS but not 100nM oAβ increased BV2 cell surface CD40 expression. **D)** Neither treatment for 24 nor 48 hrs with 100nM oAβ affected phagocytosis by BV2 cells of heat-killed E. coli bacterial particles. 50 ng/ml LPS increased phagocytosis at 48 hrs only. In all cases, data are mean ± SEM of 3-6 independent cultures, assayed in triplicate. *p<0.05 *vs*. untreated cells.

In contrast to ROS production however, 100 nM Aβ did not elicit other inflammatory changes in BV2 cells, whether assessed through production of the major inflammatory cytokine TNFα (Figure 1B), induction of the inflammatory surface phenotypic marker CD40 (Figure 1C) or phagocytosis of labelled *E. coli* bioparticles (Figure 1D). This was in marked contrast to the effects of bacterial lipopolysaccharide (LPS) which was able to evoke a clear inflammatory response from BV2 cells (Figure 1B-D).

### oAβ induces ROS production through NADPH oxidase activation, a response reversed by Fpr2/3 agonist treatment

Microglial ROS production via the enzyme NADPH oxidase, also termed NOX2, is a key response to inflammatory stimuli, primarily serving as an antimicrobial defence mechanism [31]. There is evidence for activation of this enzyme in AD [32], hence we investigated whether this was also the cellular source of ROS in our model. ROS production induced by stimulation with 100 nM oAβ was sensitive to inclusion of two different NADPH oxidase inhibitors, 1 μM diphenylene iodonium and 1 μg/ml apocynin (Figure 2A-B), strongly suggesting the involvement of this enzyme. NADPH oxidase is not the only potential cellular source of ROS however, with mitochondrial superoxide production playing a significant part in many physiological and pathological processes [33]. However, examination of BV2 cells stimulated with 100 nM oAβ found no change in mitochondrial superoxide production over 1 hr (Supplemental Figure 2A), whereas exposure to the mitochondrial complex I inhibitor rotenone (1 μM) resulted in a clear increase mitochondrial superoxide production compared to untreated (Supplemental Figure 2A-2B).

**Figure 2:**
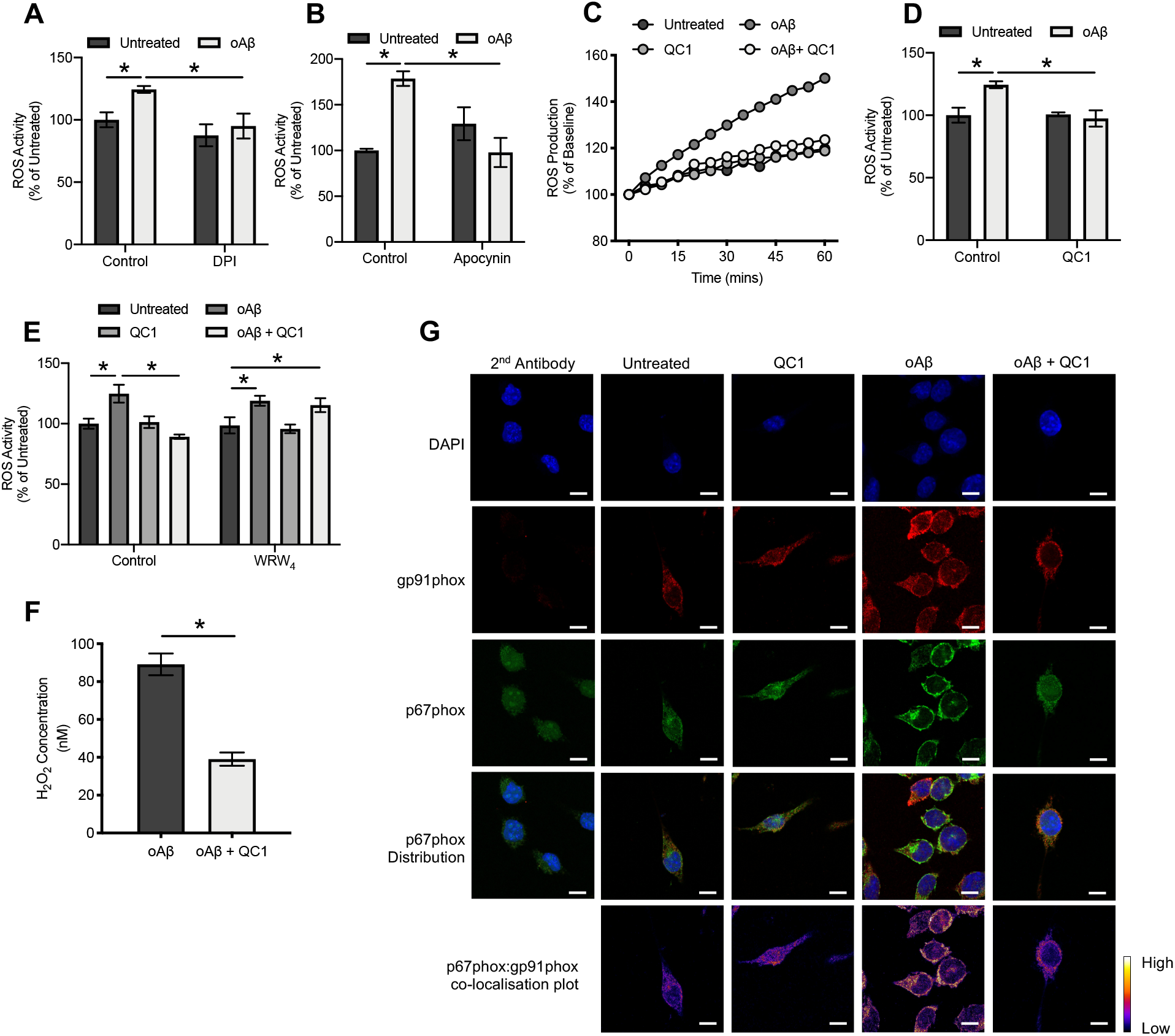
oAβ induced ROS production follows activation of NADPH oxidase and is reversed by subsequent Fpr2/3 stimulation. **A, B)** oAβ induced ROS production was prevented by 10 min pre-treatment with the NADPH oxidase inhibitors DPI (1 μM, A) and apocynin (1 μg/ml, B). **C)** Representative time-course of ROS production in untreated BV2 cells, and cells exposed to 100nM oAβ with or without subsequent stimulation with 100nM QC1 (10 min post-oAβ). **D)** Average ROS production rates for BV2 cells treated with oAβ (100 nM, 1 hr) with or without subsequent stimulation with 100 nM QC1 (10 min post-oAβ). **E)** Inclusion of the selective Fpr2/3 antagonist WRW_4_ (10μM, 10 min prior to oAβ treatment) did not affect 100 nM oAβ-induced ROS production, but prevented the effects of subsequent treatment with QC1 (100 nM, 10 min post-oAβ). **F)** Treatment of BV2 cells for 30 min with 100 nM oAβ stimulated co-localisation of the NADPH oxidase subunits p67phox (green) and gp91phox (red), an effect prevented by treatment with 100 nM QC1 administered 10 min post-oAβ. Nuclei are counter-stained with DAPI (blue); p67phox and gp91phox co-localisation intensity is represented by the false-colour plots. Graphical data are mean ± SEM of 3-6 independent cultures, assayed in triplicate, *p<0.05. Images represent cells from 3 independent cultures; scale bar = 10 μm.

Having previously showing that BV2 cells express murine Fpr2/3 [28], we investigated whether activation of this receptor could reverse oAβ-induced ROS production. Treatment of cells with the Fpr2/3 specific agonist QC1 (100nM), delivered 10 min after oAβ-stimulation restored ROS production to baseline levels (Figure 2C-2D). Moreover, this effect was sensitive to pre-treatment with the Fpr2/3 specific antagonist WRW_4_ at 10 μM (Figure 2E). Notably, production of ROS in response to oAβ itself was not affected by WRW_4_ inclusion, indicating that oAβ is not in this case signalling through Fpr2/3 (Figure 2E). Confirming these data, measurement of total cellular H_2_O_2_ revealed that whilst this species was undetectable in unstimulated cells, oAβ treatment caused significant production, an effect reversed by treatment with QC1 (Figure 2F).

NADPH oxidase is a multi-subunit enzyme, with its activation requiring the translocation of a p67 subunit from the cytosol to associate with the plasma membrane-bound gp91 subunit [32]. Confocal microscopic analysis of BV2 cells stimulated with 100 nM oAβ indicated a clear appearance of co-localised p67phox and gp91phox signal at the plasma membrane of the cells, an effect that was again prevented by subsequent treatment (10 min post-oAβ) with 100 nM QC1 (Figure 2G).

### Fpr2/3 stimulation does not modify major cellular antioxidant systems

Whilst we have shown the Fpr2/3 agonist QC1 to reverse oAβ-induced NADPH oxidase activation and ROS production, it is plausible that this could also be achieved through activation of intracellular antioxidant systems. However, neither the ratio of reduced to oxidised glutathione, nor expression of the antioxidant enzymes haem oxygenase-1 or superoxide dismutase-2 were affected by either treatment with 100nM oAβ, 100nM QC1 or a combination of the two (Figure 3). These data suggest that the ROS production-suppressing actions of Fpr2/3 activation occur through modulation at source rather than stimulation of defensive systems.

**Figure 3:**
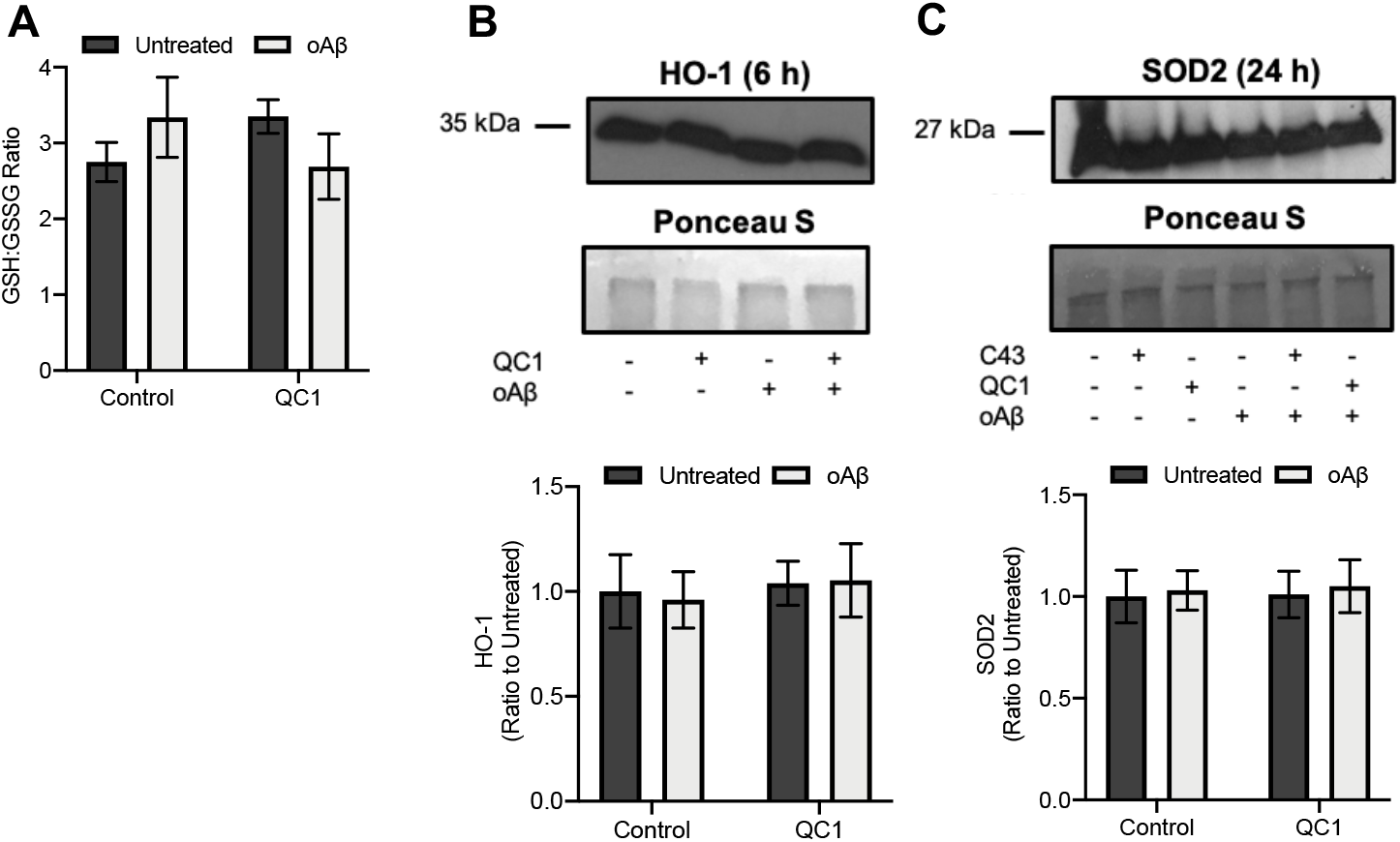
Neither treatment with oAβ nor QC1 affected major cellular antioxidant systems. **A)** The ratio of reduced (GSH) to oxidised (GSSG) glutathione within BV2 cell cytoplasm was not affected by either oAβ (100 nM, 2 hrs) or QC1 (100 nM, 10 min post-oAβ) administration. **B)** Expression of the antioxidant enzyme haem oxygenase-1 (HO-1) was not affected by treatment with oAβ (100 nM, 6 hrs) or QC1 (100 nM, 10 min post-oAβ). Sample loading was normalised to Ponceau S-defined total protein content; densitometric analysis data are mean ± SEM of 3 independent cultures. **C)** Expression of the antioxidant enzyme superoxide dismutase-2 (SOD-2) was not affected by treatment with oAβ (100 nM, 24 hrs) or QC1 (100nM, 10 min post-oAβ). All western blot analyses are representative of 3 independent cultures, with sample loading normalised to Ponceau S-defined total protein content; densitometric analysis data are mean ± SEM of 3 independent cultures, quantified in triplicate.

### Promotion of the pentose phosphate pathway by oAβ is reversed by Fpr2/3 stimulation

An important aspect of immune cell activation is a change in their preferred source of metabolic energy, with inflammatory cells tending to favour glycolysis over mitochondrial oxidative phosphorylation as their primary energy source [34]. We therefore investigated how oAβ treatment of BV2 cells would affect their metabolism through use of the Agilent Seahorse XF Analyser. Stimulation of BV2 cells with 100 nM oAβ significantly supressed basal respiration without affecting either maximal respiration or spare respiratory capacity, an effect reversed by treatment with 100 nM QC1 1 hr post-oAβ challenge (Figure 4A-E). This change in respiration resulted in a decrease in ATP production from both oxidative phosphorylation (Figure 4F) and glycolysis (Figure 4G) upon oAβ stimulation, an action again reversed by Fpr2/3 activation with QC1 (Figure 4F-4G).

**Figure 4:**
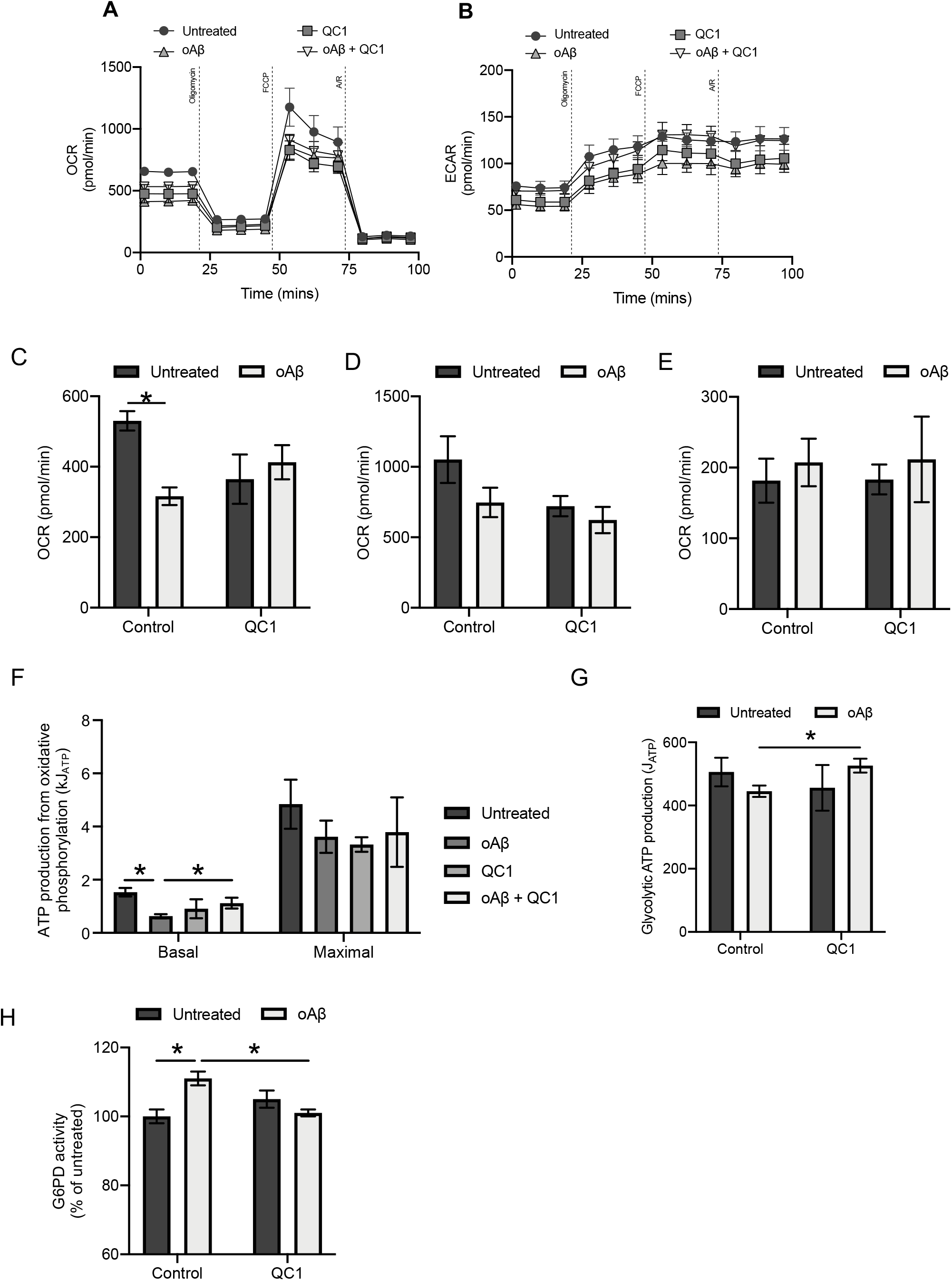
Treatment with oAβ suppresses mitochondrial respiration and promotes activity of the pentose phosphate pathway, effects reversed by subsequent activation of Fpr2/3. **A)** Typical oxygen consumption rates of untreated BV2 cells and cells treated for 24 hrs with 100 nM oAβ with or without subsequent stimulation with QC1 (100 nM, 1 hr post-oAβ), administration times for oligomycin (4 μM), FCCP (0.6 μM) and rotenone with antimycin A (both 1 μM) are indicated. **B)** Typical extracellular acidification rates for untreated BV2 cells and cells treated for 24 hrs with 100 nM oAβ with or without subsequent stimulation with QC1 (100 nM, 1 hr post-oAβ), administration times for oligomycin (4 μM), FCCP (0.6 μM) and rotenone with antimycin A (both 1 μM) are indicated. **C-E)** Treatment with 100 nM oAβ for 24 hrs significantly suppressed basal metabolic rate (C), an effect that no longer reached statistical significance after QC1 treatment (100 nM, 1 hr post-oAβ). In contrast, neither oAβ nor QC1 treatment affected maximal respiration (D) or spare respiratory capacity (E). **F)** Treatment with oAβ (100 nM, 24 hrs) significantly suppressed basal, but not maximal, ATP production due to mitochondrial oxidative phosphorylation, an effect reversed by subsequent treatment with QC1 (100 nM, 1 hr post-oAβ). **G)** ATP generation from glycolysis was unaffected by either oAβ or QC1 treatment. **H)** Activity of the rate-determining enzyme of the pentose phosphate pathway, glucose-6-phosphate dehydrogenase (G6PD) was significantly increased by treatment with 100nM oAβ (24 hrs), an effect reversed by subsequent stimulation with 100 nM QC1 (1 hr post-oAβ). All data are mean ± SEM for 3-5 independent cultures, assayed in triplicate, *p<0.05.

Production of ROS from NADPH oxidase is ultimately dependent, as its name suggests, upon a constant source of intracellular NADPH [32]. The major source of NADPH production in the cell is the pentose phosphate pathway, which siphons glucose-6-phosphate from glycolysis into the production of 6-phosphogluconate and then ribose-5-phosphate, generating NADPH in both steps [35]. As both glycolytic and mitochondrial respiratory rates were suppressed by oAβ, we investigated whether pentose phosphate pathway activity had concomitantly risen through measurement of the activity of the rate limiting enzyme for this pathway, glucose-6- phosphate dehydrogenase (G6PD). Treatment of BV2 cells with 100 nM oAβ for 24 hrs caused a significant increase in G6PD activity, an effect that was reversed to baseline upon subsequent treatment with 100nM QC1 (Figure 4H, confirming the importance of this pentose phosphate pathway shunt in the response to oAβ.

### oAβ stimulated ROS production is responsible for microglial-mediated neuronal toxicity and can be reversed by Fpr2/3 activation

Production of ROS by immune cells is primarily for the purpose of killing invading pathogens. In the context of AD however, where no infectious agent has been discovered, production of ROS may well damage bystander neurones, contributing to neurodegeneration. To investigate the relationship between oAβ-triggered microglial ROS production and neuronal health, we employed an *in vitro* co-culture model using BV2 cells and trans-retinoic acid-differentiated SH-SY5Y cells. Initial experiments revealed that 100 nM oAβ showed no direct toxicity to differentiated SH-SY5Y cells even after exposure for 24 hrs (Figure 5A). However, administration of oAβ to co-cultures significantly and selectively enhanced apoptosis of SH-SY5Y cells (Figure 5B-C) without affecting BV2 cell survival (Figure 5C), an effect that was notably prevented by treatment with 100 nM QC1 1 hr after oAβ exposure. Notably, differentiated SH-SY5Y cells did not express Fpr2/3 (Supplemental Figure 3). Confirming that either direct contact or short-lived secretory factors were responsible for BV2 cell-mediated toxicity, apoptosis of SH-SY5Y cells was not induced following treatment with conditioned medium from oAβ-stimulated BV2 cultures (Figure 5D). Finally, to test whether BV2 cell ROS production was the mediating agent for SH-SY5Y cytotoxicity, the experiment was repeated in the presence of the antioxidant molecule α-tocopherol (10 μM). Inclusion of this antioxidant prevented SH-SY5Y apoptosis in co-cultures treated with oAβ, indicating a direct mediatory role of microglial ROS production.

**Figure 5:**
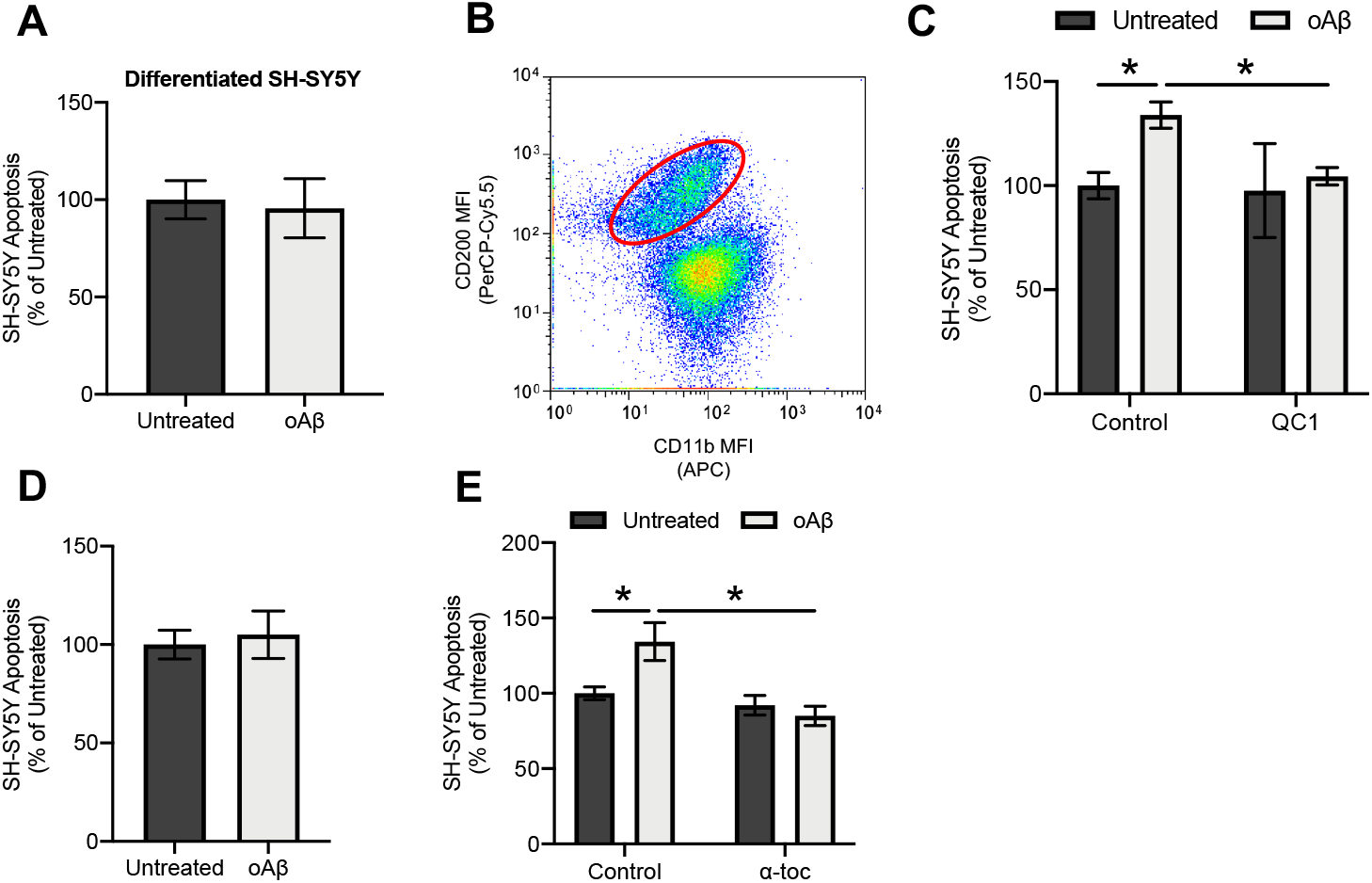
Treatment with oAβ induces differentiated SH-SY5Y neuronal apoptosis only in the presence of microglia, acting through Fpr2/3 sensitive ROS release. **A)** Treatment with oAβ (100 nM, 48 hrs) had no effect on *trans*-retinoic acid (tRA)-differentiated SH-SY5Y cell viability; data are mean ± SEM of 5 independent cultures, assayed in triplicate. **B)** Separation of tRA-differentiated SH-SY5Y neurons from BV2 cells grown in co-culture on the basis of differential CD200 and CD11b expression, plot is representative of 3 independent cultures. **C)** Treatment of co-cultures of BV2 and tRA-differentiated SH-SY5Y neurons with oAβ (100 nM, 48 hrs) induces significant SH-SY5Y apoptosis, an effect prevented by subsequent treatment with QC1 (100 nM, 10 min post-oAβ). **D)** Conditioned medium from BV2 cells treated or not with 100nM oAβ (24 hrs) had no effect on tRA-differentiated SH-SY5Y neuronal apoptosis following exposure for 48 hrs. **E)** Inclusion of the antioxidant α-tocopherol (10 μM) in co-cultures of BV2 cells and tRA-differentiated SH-SY5Y neurons prevented oAβ- induced (100 nM, 48 hrs) neuronal apoptosis; data are mean ± SEM for 3-6 independent cultures, assayed in triplicate, *p<0.05.

## Discussion

Despite over 300 clinical trials having been performed targeting either of the proposed toxic mediators in AD, Aβ and hyperphosphorylated tau, we do not as yet have any successful therapeutic approaches for the disease. This suggests that, at the least, these two proteins cannot be the sole factors driving the disease [36]. Increasingly, the role of neuroinflammation and the behaviour of microglia in AD has come under investigation [3,37], an approach given further impetus by reports that ablation of microglia can halt brain atrophy in murine models of Aβ-driven disease [38] and tauopathy [39]. Microglial activation can be both beneficial and damaging, hence strategies that can control excessive inflammatory activity and promote a pro-resolving phenotype may be of great potential for therapeutic use. In this study, we have used an *in vitro* cellular model to provide proof-of-principle evidence for the targeting of the pro-resolving receptor Fpr2/3 as a mechanism to restrain microglial behaviour and limit the ability of these cells to damage bystander neurones.

Here, we report that alongside its well-characterised function in resolving inflammation and efferocytosis [40], Fpr2/3 activation can reverse oAβ-induced ROS production through deactivation of NADPH oxidase activity. Activation of microglial NADPH oxidase by oAβ is well supported [41,42], and may be critical in triggering neuroinflammation, given the damaging effects of oxidative stress for neurones, as has been reported in traumatic brain injury [43]. Future work will determine whether the *in vitro* findings we report here can be extended to the *in vivo* situation, but if so, they suggest that the use of Fpr2/3 agonists capable of reversing NADPH oxidase activation may be of therapeutic potential for AD.

The effects of Fpr2/3 activation upon oAβ-induced ROS production are mirrored by changes in microglial metabolic phenotype. The importance of cellular metabolism in regulating immune cell phenotype has become increasingly evident over the past few years, with a shift from mitochondrial respiration to a glycolysis-dominant metabolism being closely associated with a pro-inflammatory phenotype [34]. Metabolic changes in AD are well supported [44], but the relationship between these changes and disease pathology is unclear. In the current study, exposure of microglia to oAβ suppressed mitochondrial respiration, but rather than being accompanied by changes to glycolysis, it was associated with a significant diversion of glucose to the pentose phosphate pathway. Presumably, this was due to increased NADPH demand associated with NADPH oxidase-driven ROS production, as seen in peripheral macrophages [45]. Notably, Fpr2/3 agonist treatment was able to reverse the effects of oAβ on microglial metabolism, targeting both the pentose phosphate pathway and the mitochondria. This data adds to the increasing evidence suggesting that Fpr2/3 not only suppresses pro-inflammatory mediator production [16,46], but aids in the regulation of the underlying metabolic changes that occur in activated immune cells, as we have recently shown in peripheral macrophages [14]. Importantly, microglia rapidly upregulate Fpr2/3 expression following inflammatory insult [47], and whilst the effects of its stimulation on neuroinflammation can be agonist dependent [48], selective Fpr2/3 activation contributes to neuroinflammatory resolution in a murine model of AD [46].

However, a notable finding of the current work is that we were unable to detect evidence for an oAβ-induced microglial inflammatory response, in contrast to previous *in vitro* studies [24,47,51]. This does not appear to be due to deficiency in the BV2 cells themselves, as stimulation with bacterial lipopolysaccharide was still able to trigger a potent inflammatory response. Notably, studies that report pro-inflammatory effects of oAβ have commonly used micromolar concentrations of the peptide, several orders of magnitude greater than levels reported to occur in the human brain in AD [30]. This suggests that the direct pro-inflammatory effects of oAβ seen in *in vitro* may not be fully recapitulated *in vivo*, although this requires further validation. Nevertheless, oAβ was clearly able to induce ROS production at levels capable of damaging bystander cells, which if replicated *in vivo* may be a driving factor in ongoing neuronal damage and secondary neuroinflammation seen in AD. Production of ROS is far from the sole damaging effect of oAβ in the brain, as is borne out by the, at best, equivocal results from clinical trials of antioxidants in AD [50,51]. Nevertheless, targeting a receptor with the potential to suppress ROS production, restore microglial metabolic homeostasis and promote resolution, as is the case for Fpr2/3, has significant potential for therapeutic development.

The goal of this study was to provide proof-of-principle evidence that exploitation of human FPR2 may hold promise as a therapeutic target for AD research. Evidently there are limitations in how far the current study should be interpreted, with a particular need for further *in vivo* validation of our findings. Nonetheless, the data presented do suggest that this receptor would be a suitable target for anti-oxidative and metabolic therapeutic development for further AD research, particularly as the effects of Fpr2/3 stimulation were apparent when stimulation occurred after oAβ treatment. This study therefore adds novel insights into the role of Fpr2/3 in modulating microglial oxidative stress and metabolism, holding promise for select Fpr2/3 agonists as potent and effective treatment options for inflammatory disease [9].

## Conclusions

This study has identified that although pathologically relevant concentrations of oAβ do not appear to directly stimulate an inflammatory microglial phenotype, they are potent activators of microglial ROS production via NADPH oxidase, and consequent changes to metabolic phenotype. Moreover, activation of the pro-resolving receptor Fpr2/3 was able to reverse these oAβ-induced changes and protect bystander neurones from damage. These data suggest that manipulation of Fpr2/3 may be an important target for future therapeutic development in neuroinflammatory conditions such as Alzheimer’s disease.

## List of abbreviations

AD: Alzheimer’s disease
FPR2: Human formyl peptide receptor 2
Fpr2/3: Murine formyl peptide receptors 2/3
GSH: Reduced glutathione
GSSG: Oxidised glutathione
oAβ: Oligomeric β-amyloid 1-42
ROS: Reactive oxygen species
TNFα: Tumour necrosis factor alpha
WRW_4_: Trp-Arg-Trp-Trp-Trp-Trp

---

**Supplementary Figure 1:**
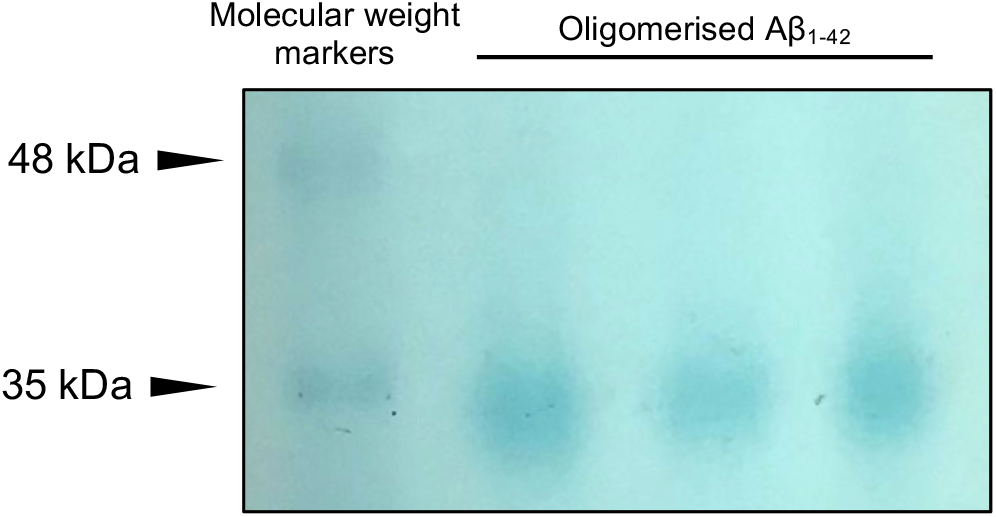
Approximate molecular weight of Aβ1-42 oligomers following polyacrylamide gel electrophoresis under non-denaturing conditions. As monomeric Aβ_1-42_ has a molecular weight of 4.51 kDa, the apparent molecular weight of approximately 35kDa suggest that oAβ species were hexamers/heptamers.

**Supplementary Figure 2:**
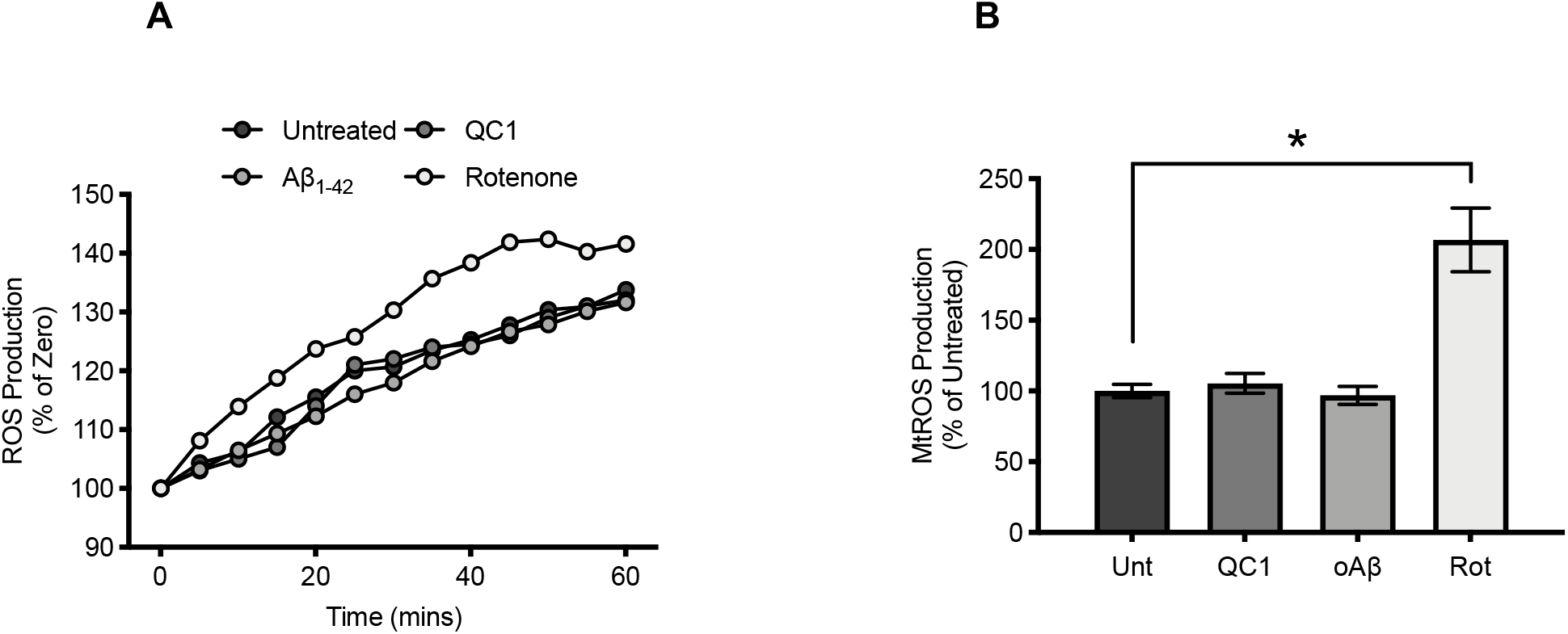
Mitochondrial ROS production is not stimulated in microglia by oAβ treatment. **A)** Representative time-course of mitochondrial superoxide production in untreated BV2 cells and cells exposed to 100 nM oAβ, 100nM QC1 or 1 μM rotenone for 1 hr. **B)** Average mitochondrial superoxide production rates for untreated BV2 cells and cells treated with oAβ (100 nM, 1 hr), QC1 (100 nM, 1 hr) or rotenone (ROT; 1 μM, 1 hr); data are mean ± SEM of 4 independent cultures, assayed in triplicate, *p<0.05.

**Supplementary Figure 3:**
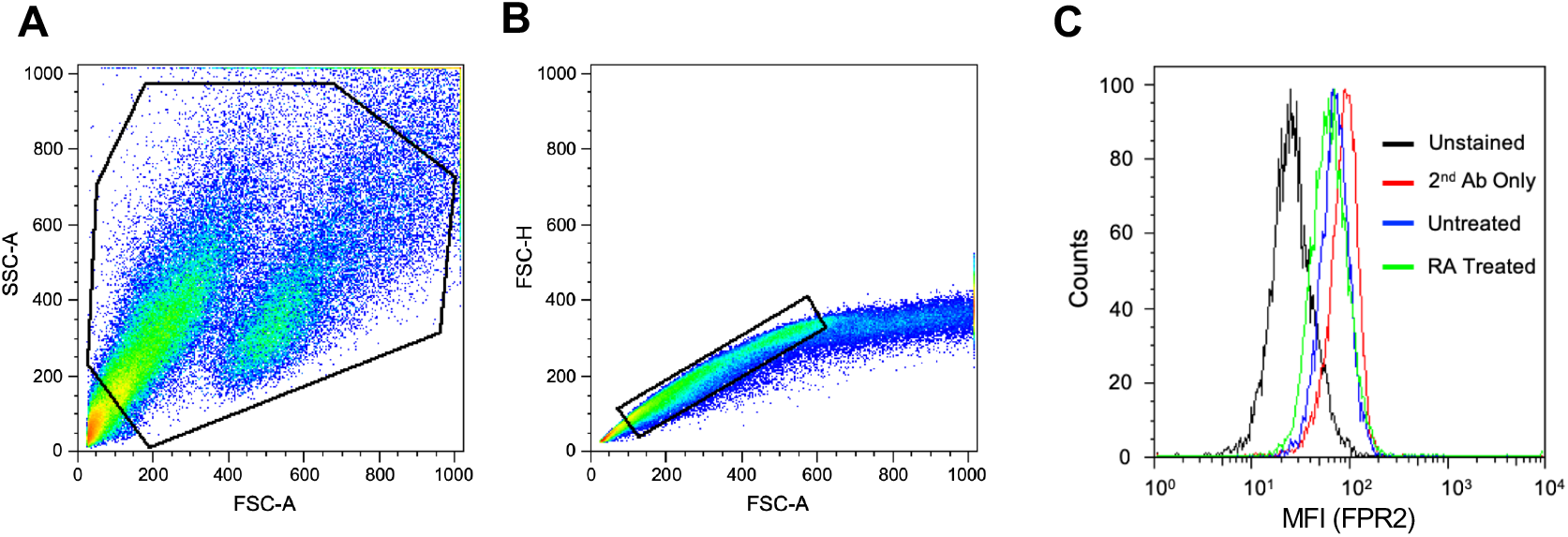
FPR2 is not expressed by SH-SY5Y cells. **A)** representative forward scatter-side scatter histogram plot and gating strategy. **B**) representative histogram plot to exclude cell doublets. **C**) histogram of relative FPR2 staining. Neither naïve or transretinoic acid induced differentiated SH-SY5Y cells expressed FPR2.

## References

1. United Nations. World Population Prospects: The 2015 Revision, Key Findings and Advance Tables. World Popul. Prospect. 2015.

2. Prince MJ, Wimo A, Guerchet MMM, Gemma-Claire A, Wu Y-T, Prina M, et al. World Alzheimer Report 2015 - The Global Impact of Dementia: An analysis of prevalence, incidence, cost and trends. Alzheimer’s Dis. Int. Alzheimer’s Disease International; 2015.

3. Heneka MT, Carson MJ, Khoury J El, Landreth GE, Brosseron F, Feinstein DL, et al. Neuroinflammation in Alzheimer’s disease. Lancet Neurol. 2015;14:388–405.

4. Ransohoff RM, El Khoury J. Microglia in Health and Disease. Cold Spring Harb. Perspect. Cold Spring Harbor Laboratory Press; 2015;8:a020560.

5. Taipa R, Ferreira V, Brochado P, Robinson A, Reis I, Marques F, et al. Inflammatory pathology markers (activated microglia and reactive astrocytes) in early and late onset Alzheimer disease: a *post mortem* study. Neuropathol. Appl. Neurobiol. 2018;44:298–313.

6. Spangenberg EE, Green KN. Inflammation in Alzheimer’s disease: Lessons learned from microglia-depletion models. Brain. Behav. Immun. 2017;61:1–11.

7. Heneka MT, Kummer MP, Latz E. Innate immune activation in neurodegenerative disease. Nat. Publ. Gr. 2014;14.

8. Yu Y, Ye RD. Microglial Aβ Receptors in Alzheimer’s Disease. Cell. Mol. Neurobiol. 2014;35:71–83.

9. Sugimoto MA, Vago JP, Perretti M, Teixeira MM. Mediators of the Resolution of the Inflammatory Response. Trends Immunol. Elsevier Ltd; 2019;40:212–27.

10. Dufton N, Hannon R, Brancaleone V, Dalli J, Patel HB, Gray M, et al. Anti-inflammatory role of the murine formyl-peptide receptor 2: ligand-specific effects on leukocyte responses and experimental inflammation. J. Immunol. 2010;184:2611–9.

11. Solito E, Kamal A, Russo-Marie F, Buckingham JC, Marullo S, Perretti M. A novel calcium-dependent proapoptotic effect of annexin 1 on human neutrophils. FASEB J. 2003;17:1544–6.

12. Drechsler M, de Jong R, Rossaint J, Viola JR, Leoni G, Wang JM, et al. Annexin A1 counteracts chemokine-induced arterial myeloid cell recruitment. Circ. Res. 2015;116:827–35.

13. McArthur S, Gobbetti T, Kusters DHM, Reutelingsperger CP, Flower RJ, Perretti M. Definition of a Novel Pathway Centered on Lysophosphatidic Acid To Recruit Monocytes during the Resolution Phase of Tissue Inflammation. J. Immunol. 2015;195:1500733.

14. McArthur S, Juban G, Gobbetti T, Desgeorges T, Theret M, Gondin J, et al. Annexin A1 drives macrophage skewing to accelerate muscle regeneration through AMPK activation. J. Clin. Invest. 2020;124635.

15. Scannell M, Flanagan MB, deStefani A, Wynne KJ, Cagney G, Godson C, et al. Annexin-1 and peptide derivatives are released by apoptotic cells and stimulate phagocytosis of apoptotic neutrophils by macrophages. J. Immunol. 2007;178:4595–605.

16. Gobbetti T, Coldewey SMSM, Chen J, McArthur S, le Faouder P, Cenac N, et al. Nonredundant protective properties of FPR2/ALX in polymicrobial murine sepsis. Proc. Natl. Acad. Sci. U. S. A. 2014;111:18685–90.

17. Kain V, Jadapalli JK, Tourki B, Halade G V. Inhibition of FPR2 impaired leukocytes recruitment and elicited non-resolving inflammation in acute heart failure. Pharmacol. Res. 2019;146:104295.

18. Petri MH, Laguna-Fernandez A, Arnardottir H, Wheelock CE, Perretti M, Hansson GK, et al. Aspirin-triggered lipoxin A4 inhibits atherosclerosis progression in apolipoprotein E ^−/−^ mice. Br. J. Pharmacol. 2017;

19. Ho CF-Y, Ismail NB, Koh JK-Z, Gunaseelan S, Low Y-H, Ng Y-K, et al. Localisation of Formyl-Peptide Receptor 2 in the Rat Central Nervous System and Its Role in Axonal and Dendritic Outgrowth. Neurochem. Res. 2018;43:1587–98.

20. Iribarren P, Zhou Y, Hu J, Le Y, Wang JM. Role of formyl peptide receptor-like 1 (FPRL1/FPR2) in mononuclear phagocyte responses in Alzheimer disease. Immunol. Res. 2005;31:165–76.

21. Cui YH, Le Y, Zhang X, Gong W, Abe K, Sun R, et al. Up-regulation of FPR2, a chemotactic receptor for amyloid beta 1-42 (A beta 42), in murine microglial cells by TNF alpha. Neurobiol. Dis. 2002;10:366–77.

22. Le Y, Gong W, Tiffany HL, Tumanov A, Nedospasov S, Shen W, et al. Amyloid (beta)42 activates a G-protein-coupled chemoattractant receptor, FPR-like-1. J. Neurosci. 2001;21:RC123.

23. Tiffany HL, Lavigne MC, Cui YH, Wang JM, Leto TL, Gao JL, et al. Amyloid-beta induces chemotaxis and oxidant stress by acting at formylpeptide receptor 2, a G protein-coupled receptor expressed in phagocytes and brain. J. Biol. Chem. American Society for Biochemistry and Molecular Biology; 2001;276:23645–52.

24. Ries M, Loiola R, Shah UN, Gentleman SM, Solito E, Sastre M. The anti-inflammatory Annexin A1 induces the clearance and degradation of the amyloid-β peptide. J. Neuroinflammation. 2016;13:234.

25. Stine B, Jungbauer L, Chunjiang Y, LaDu M. Preparing Synthetic Aβ in Different Aggregation States. Methods Mol. Biol. 2011;670:13–32.

26. Shipley MM, Mangold CA, Szpara ML. Differentiation of the SH-SY5Y Human Neuroblastoma Cell Line. J. Vis. Exp. MyJoVE Corporation; 2016;53193.

27. Mookerjee SA, Gerencser AA, Nicholls DG, Brand MD. Quantifying intracellular rates of glycolytic and oxidative ATP production and consumption using extracellular flux measurements. J. Biol. Chem. American Society for Biochemistry and Molecular Biology; 2017;292:7189–207.

28. McArthur S, Cristante E, Paterno M, Christian H, Roncaroli F, Gillies GEE, et al. Annexin A1: a central player in the anti-inflammatory and neuroprotective role of microglia. J Immunol. 2010/10/22. 2010;185:6317–28.

29. Loiola RA, Wickstead ES, Solito E, McArthur S. Estrogen promotes pro-resolving microglial behaviour and phagocytic cell clearance through the actions of annexin A1. Front. Endocrinol. (Lausanne). Cold Spring Harbor Laboratory; 2019;10:420.

30. Helmond Z Van, Miners JS, Kehoe PG, Love S. Higher Soluble Amyloid b Concentration in Frontal Cortex of Young Adults than in Normal Elderly or Alzheimer’s Disease. 2010;20:787–93.

31. Spooner R, Yilmaz O. The role of reactive-oxygen-species in microbial persistence and inflammation. Int. J. Mol. Sci. Multidisciplinary Digital Publishing Institute (MDPI); 2011;12:334–52.

32. Ma MW, Wang J, Zhang Q, Wang R, Dhandapani KM, Vadlamudi RK, et al. NADPH oxidase in brain injury and neurodegenerative disorders. Mol. Neurodegener. 2017;12:7.

33. Zorov DB, Juhaszova M, Sollott SJ. Mitochondrial reactive oxygen species (ROS) and ROS-induced ROS release. Physiol. Rev. American Physiological Society; 2014;94:909–50.

34. O’Neill LAJ, Kishton RJ, Rathmell J. A guide to immunometabolism for immunologists. Nat. Rev. Immunol. 2016;16:553–65.

35. Grant CM. Metabolic reconfiguration is a regulated response to oxidative stress. J. Biol. BioMed Central; 2008. p. 1.

36. Makin S. The amyloid hypothesis on trial. Nature. 2018;559:S4–7.

37. Merlini M, Rafalski VA, Rios Coronado PE, Gill TM, Ellisman M, Muthukumar G, et al. Fibrinogen Induces Microglia-Mediated Spine Elimination and Cognitive Impairment in an Alzheimer’s Disease Model. Neuron. 2019;101:1099–1108.e6.

38. Spangenberg EE, Lee RJ, Najafi ARR, Rice RAA, Elmore MRPRP, Blurton-Jones M, et al. Eliminating microglia in Alzheimer’s mice prevents neuronal loss without modulating amyloid-β pathology. Brain. Oxford University Press; 2016;139:1265–81.

39. Shi Y, Manis M, Long J, Wang K, Sullivan PM, Remolina Serrano J, et al. Microglia drive APOE-dependent neurodegeneration in a tauopathy mouse model. J. Exp. Med. Rockefeller University Press; 2019;jem.20190980.

40. Corminboeuf O, Leroy X. FPR2/ALXR Agonists and the Resolution of Inflammation. J. Med. Chem. 2015;58:537–59.

41. Abubaker AA, Vara D, Visconte C, Eggleston I, Torti M, Canobbio I, et al. Amyloid Peptide β1-42 Induces Integrin αIIbβ3 Activation, Platelet Adhesion, and Thrombus Formation in a NADPH Oxidase-Dependent Manner. Oxid. Med. Cell. Longev. 2019;2019:1050476.

42. Alawieyah Syed Mortadza S, Sim JA, Neubrand VE, Jiang L-H. A critical role of TRPM2 channel in Aβ42-induced microglial activation and generation of tumor necrosis factor-α. Glia. 2018;66:562–75.

43. Kumar A, Barrett JP, Alvarez-Croda D-M, Stoica BA, Faden AI, Loane DJ. NOX2 drives M1-like microglial/macrophage activation and neurodegeneration following experimental traumatic brain injury. Brain. Behav. Immun. 2016;58:291–309.

44. Mostafavi S, Gaiteri C, Sullivan SE, White CC, Tasaki S, Xu J, et al. A molecular network of the aging human brain provides insights into the pathology and cognitive decline of Alzheimer’s disease. Nat. Neurosci. 2018;21:811–9.

45. Nagy C, Haschemi A. Time and Demand are Two Critical Dimensions of Immunometabolism: The Process of Macrophage Activation and the Pentose Phosphate Pathway. Front. Immunol. Frontiers; 2015;6:164.

46. Kantarci A, Aytan N, Palaska I, Stephens D, Crabtree L, Benincasa C, et al. Combined administration of resolvin E1 and lipoxin A4 resolves inflammation in a murine model of Alzheimer’s disease. Exp. Neurol. 2018;300:111–20.

47. Chen K, Iribarren P, Huang J, Zhang L, Gong W, Cho EH, et al. Induction of the Formyl Peptide Receptor 2 in Microglia by IFN- and Synergy with CD40 Ligand. J. Immunol. 2007;178:1759–66.

48. He H-Q, Ye R. The Formyl Peptide Receptors: Diversity of Ligands and Mechanism for Recognition. Molecules. Multidisciplinary Digital Publishing Institute; 2017;22:455.

49. Liu H, Wang JJ, Wang JJ, Wang P, Xue Y. Paeoniflorin attenuates Aβ1-42-induced inflammation and chemotaxis of microglia in vitro and inhibits NF-κB- and VEGF/Flt-1 signaling pathways. Brain Res. 2015;1618:149–58.

50. D’Cunha NM, Georgousopoulou EN, Dadigamuwage L, Kellett J, Panagiotakos DB, Thomas J, et al. Effect of long-term nutraceutical and dietary supplement use on cognition in the elderly: a 10-year systematic review of randomised controlled trials. Br. J. Nutr. 2018;119:280–98.

51. McCleery J, Abraham RP, Denton DA, Rutjes AW, Chong L-Y, Al-Assaf AS, et al. Vitamin and mineral supplementation for preventing dementia or delaying cognitive decline in people with mild cognitive impairment. Cochrane Database Syst. Rev. 2018;11:CD011905.

